# A Prokaryotic Membrane Sculpting BAR Domain Protein

**DOI:** 10.1101/2020.01.30.926147

**Authors:** Daniel A. Phillips, Lori A. Zacharoff, Cheri M. Hampton, Grace W. Chong, Anthony P. Malanoski, Lauren Ann Metskas, Shuai Xu, Lina J. Bird, Brian J. Eddie, Grant J. Jensen, Lawrence F. Drummy, Mohamed Y. El-Naggar, Sarah M. Glaven

## Abstract

Bin/Amphiphysin/RVS (BAR) domain proteins belong to a superfamily of coiled-coil proteins influencing membrane curvature in eukaryotes and are associated with vesicle biogenesis, vesicle-mediated protein trafficking, and intracellular signaling. Here we report the first prokaryotic BAR domain protein, BdpA, from *Shewanella oneidensis* MR-1, known to produce redox-active membrane vesicles and micrometer-scale outer membrane extensions (OMEs). BdpA is required for uniform size distribution of membrane vesicles and scaffolding OMEs into a consistent diameter and curvature. Cryogenic transmission electron microscopy reveals a strain lacking BdpA produces lobed, disordered OMEs rather than membrane tubes produced by the wild type strain. Overexpression of BdpA promotes OME formation during conditions where they are less common. Heterologous expression results in OME production in *Marinobacter atlanticus* and *Escherichia coli*. Based on the ability of BdpA to alter membrane curvature *in vivo*, we propose that BdpA and its homologs comprise a newly identified class of prokaryotic BAR (P-BAR) domains.

## Introduction

Eukaryotic Bin/Amphiphysin/Rvs (BAR) domain-containing proteins generate membrane curvature through electrostatic interactions between positively charged amino acids and negatively charged lipids, scaffolding the membrane along the intrinsically curved surface of the antiparallel coiled-coil protein dimers (1–4). Some BAR domain-containing proteins, such as the N-BAR protein BIN1, contain amphipathic helical wedges that insert into the outer membrane leaflet and can assist in membrane binding (5). Other BAR domains can be accompanied by a membrane targeting domain, such as PX for phosphoinositide binding (6, 7), in order to direct membrane curvature formation at specific sites, as is the case with sorting nexin BAR proteins (8). The extent of accumulation of BAR domain proteins at a specific site can influence the degree of the resultant membrane curvature (9), and tubulation events arise as a consequence of BAR domain multimerization in conjunction with lipid binding (10). Interactions between BAR domain proteins and membranes resolve membrane tension, promote membrane stability, and aid in localizing cellular processes, such as actin binding, signaling through small GTPases, membrane vesicle scission, and vesicular transport of proteins (11–13). Despite our knowledge of numerous eukaryotic BAR proteins spanning a variety of modes of curvature formation, membrane localizations, and subtypes (N-BAR, F-BAR, and I-BAR), characterization of a functional prokaryotic BAR domain protein has yet to be reported.

Bacterial cell membrane curvature can be observed during the formation of outer membrane vesicles (OMV) and outer membrane extensions (OME). OMV formation is ubiquitous and has many documented functions (14). OMEs are less commonly observed, remain attached to the cell, and various morphologies can be seen extending from single cells including *Myxococcus xanthus* (15, 16), flavobacterium strain Hel3_A1_48 (17), *Vibrio vulnificus* (18), *Francisella novicida* (19), *Shewanella oneidensis* (20–23), and as cell-cell connections in *Bacillus subtilis* (24–26) and *Escherichia coli* (27). Several bacterial proteins have demonstrated membrane tubule formation capabilities *in vitro* (28–33), but despite the growing number of reports, proteins involved in shaping bacterial membranes into OMV/Es have yet to be identified. Recently, researchers have begun to suspect that OMV and OME formation has some pathway overlap (17), and it is proposed that proteins are necessary to stabilize these structures (34).

*Shewanella oneidensis* is a model organism for extracellular electron transfer (EET), a mode of respiration whereby electrons traverse the inner membrane, periplasm, and outer membrane via multiheme cytochromes to reach exogenous insoluble terminal electron acceptors, such as metals and electrodes (35, 36). *S. oneidensis* is known to produce redox-active OMVs (37) and OMEs coated with mulitheme cytochromes, particularly upon surface attachment (20, 22, 37). However, little is known about their formation mechanism, control of shape or curvature, and electrochemical properties that influence EET function.

Previously, OMEs of *S. oneidensis* were shown to transition between chains of vesicles and tubules. We identified a component critical to this structural transition as the first BAR domain protein in prokaryotes, which we term BdpA (<u>B</u>AR <u>d</u>omain-like protein A). Through comparative proteomics, cryogenic electron microscopy, and molecular biology, we show that BdpA is enriched in OME/Vs, regulates size of OMVs, and controls the shape of OMEs. Likewise, BdpA confers OME formation capabilities when expressed in other bacteria, showing mechanistic evidence of BAR domain protein-mediated tubule formation *in vivo*. This study provides a framework for prokaryotic BAR domain characterization, and putative BAR domain-containing BdpA homologs in other bacteria suggest BAR domain protein-mediated membrane sculpting is an evolutionarily conserved function.

## Results and Discussion

### *S. oneidensis* OMVs are redox-active and enriched with BdpA

OMVs were purified from *S. oneidensis* cells grown in batch cultures to characterize their redox features and unique proteome, as well as to identify putative membrane shaping proteins. Cryogenic transmission electron microscopy (cryo-TEM) tomographic reconstruction slices of the purified samples showed uniform OMVs with the characteristic single membrane phenotype and an approximate diameter of 200 nm (Fig. 1a). Previous measurements showed OMVs can reduce extracellular electron acceptors (37) and that vesicles from *G. sulfurreducens* can mediate electron transfer (38). Electrochemical activity of multiheme cytochrome complex MtrCAB and their ability to mediate micrometer-scale electron transport has been characterized in whole cells (39), but no electrochemical characterization of OME/Vs has been reported that link activity to multiheme cytochromes. Here, electrochemical measurements of isolated OMVs were performed to determine if purified OMVs maintain the characteristic redox features of the MtrCAB complex when detached from cells. Cyclic voltammetry (CV) demonstrated redox activity of isolated membrane vesicles adhered to a gold electrode via self-assembled monolayers (Fig. 1b). The first derivative (Fig. 1b inset) revealed a prominent peak with a midpoint potential of 66 mV and a smaller peak at −25 mV versus a standard hydrogen reference electrode (SHE). This midpoint potential is consistent with the characteristics of multiheme cytochromes such as MtrC/OmcA from previous microbial electrochemical studies (39, 40), suggesting that the extracellular redox molecules of the cellular outer membrane extends to OMVs.

**Fig 1.**
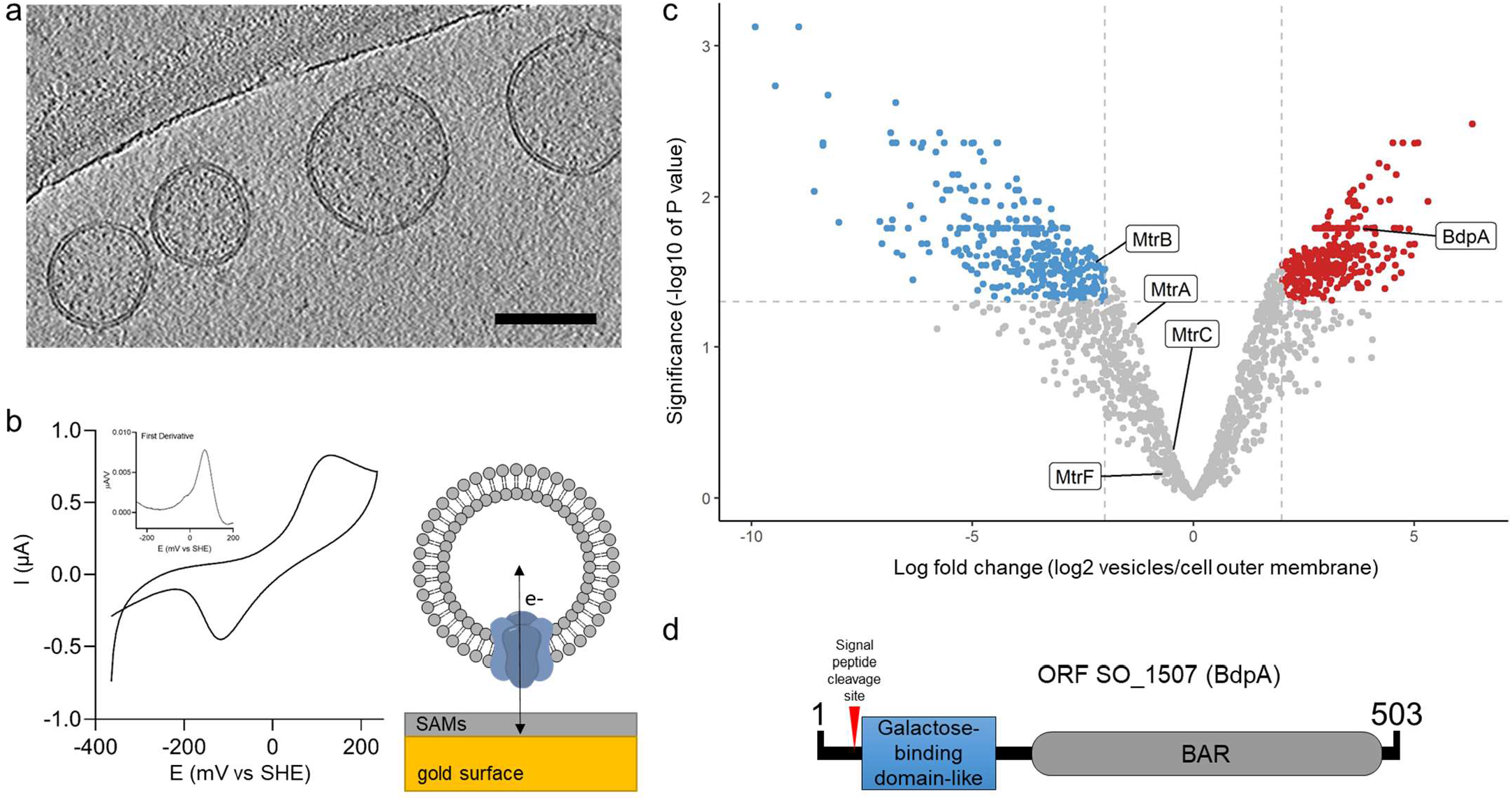
Redox active vesicles are enriched with BAR domain protein BdpA. **a**. Cryoelectron tomography image of *S. oneidensis* MR-1 outer membrane vesicles (OMVs) (scale = 200 nm). **b**. Cyclic voltammetry of vesicles adhered to gold electrode via small self-assembled monolayers, as diagramed. Inset shows first derivative of anodic scan. **c**. Volcano plot of vesicle proteome compared to cell-associated outer membrane (red = enriched in vesicles, blue = enriched in cell-associated outer membrane). **d**. Schematic of BdpA domains.

The proteome of the OMVs was compared to the proteome of purified outer membranes extracted from whole cells. Using a label-free quantification method (41), significant differences in the ratio of individual proteins in the vesicle to the outer membrane could be computed (log fold change) (Fig 1c). The proteome of the purified OMVs showed 328 proteins were significantly enriched in the vesicles as compared to the outer membrane, and 314 proteins were significantly excluded from the vesicles (Fig. 1c, Supplemental table 1). MtrCAB cytochromes were present in the OMVs as well as the outer membrane, supporting redox activity of OMVs observed by CV. Active protein sorting into eukaryotic vesicles is a coordinated process involving protein sorting signal recognition, localized membrane protein recruitment, initiation of membrane curvature induction, and coating nascent vesicles with protein scaffolds (42). Several proteins significantly enriched in the vesicles were identified that could contribute to OMV formation, including the murein transglycosylase, the peptidoglycan degradation enzyme holin, cell division coordinator CpoB, and a highly enriched putative BAR domain-containing protein encoded by the gene at open reading frame SO_1507, hereafter named BAR domain-like protein A (BdpA) (Fig. 1d).

Vesicle enrichment of BdpA led us to the hypothesis that BdpA could be involved in membrane shaping of OMVs based on the role of BAR domain proteins in eukaryotes. The C-terminal BAR domain of BdpA is predicted to span an alpha-helical region from AA 276-451 (E-value = 2.96e-03); however, since the identification of the protein is based on homology to the eukaryotic BAR domain consensus sequence (cd07307), it is possible that the BAR domain region extends beyond these bounds (Fig. 1d). Coiled coil prediction (43) suggests BdpA exists in an oligomeric state of antiparallel alpha-helical dimers, as is the case for all known BAR domain proteins (2, 44–46). The predicted structure of the BAR domain-containing region generated in i-TASSER (47) shows a 3 helix bundle (Supplemental Fig. 1a). When the BdpA BAR domain monomers were aligned to the dimeric structure of the F-BAR protein Hof1p (48), the predicted dimer interface residues of BdpA came into proximity, revealing an intrinsically curved dimer with positively charged residues along the concave surface (Supplemental Fig. 1b). BdpA has a N-terminal signal peptide with predicted cleavage sites between amino acids 22-23, suggesting non-cytoplasmic localization (Fig. 1d). A galactose-binding domain-like region positioned immediately downstream of the signal peptide supports lipid targeting activity seen in other BAR domain proteins, such as the eukaryotic sorting nexins (49) which have phox (PX) domains that bind phosphoinositides (50). The *S. oneidensis* rough-type lipopolysaccharide (LPS) contains 2-acetamido-2-deoxy-D-galactose (51), which suggests possible localization of the protein to the outer leaflet of the outer membrane.

### BdpA controls size distribution of vesicles

In eukaryotic cells, BAR domain proteins are implicated in vesicle formation (52, 53) and regulation of vesicle size (54). To determine whether BdpA influences vesicle morphology, OMVs were harvested from wild type (WT) cells and cells in which the gene for BdpA had been deleted (Δ*bdpA*), and their diameters were measured by dynamic light scattering (DLS). WT OMVs had a median diameter of 190 nm with little variability in the population (standard deviation (s.d.) = ±21 nm), while the diameters of Δ*bdpA* OMVs were distributed over a wider range with a median value of 280 nm (s.d. = ± 131 nm (Fig 2a). The data suggest BdpA controls vesicle diameter in membrane structures *ex vivo*, potentially acting by stabilizing OMVs.

**Fig 2.**
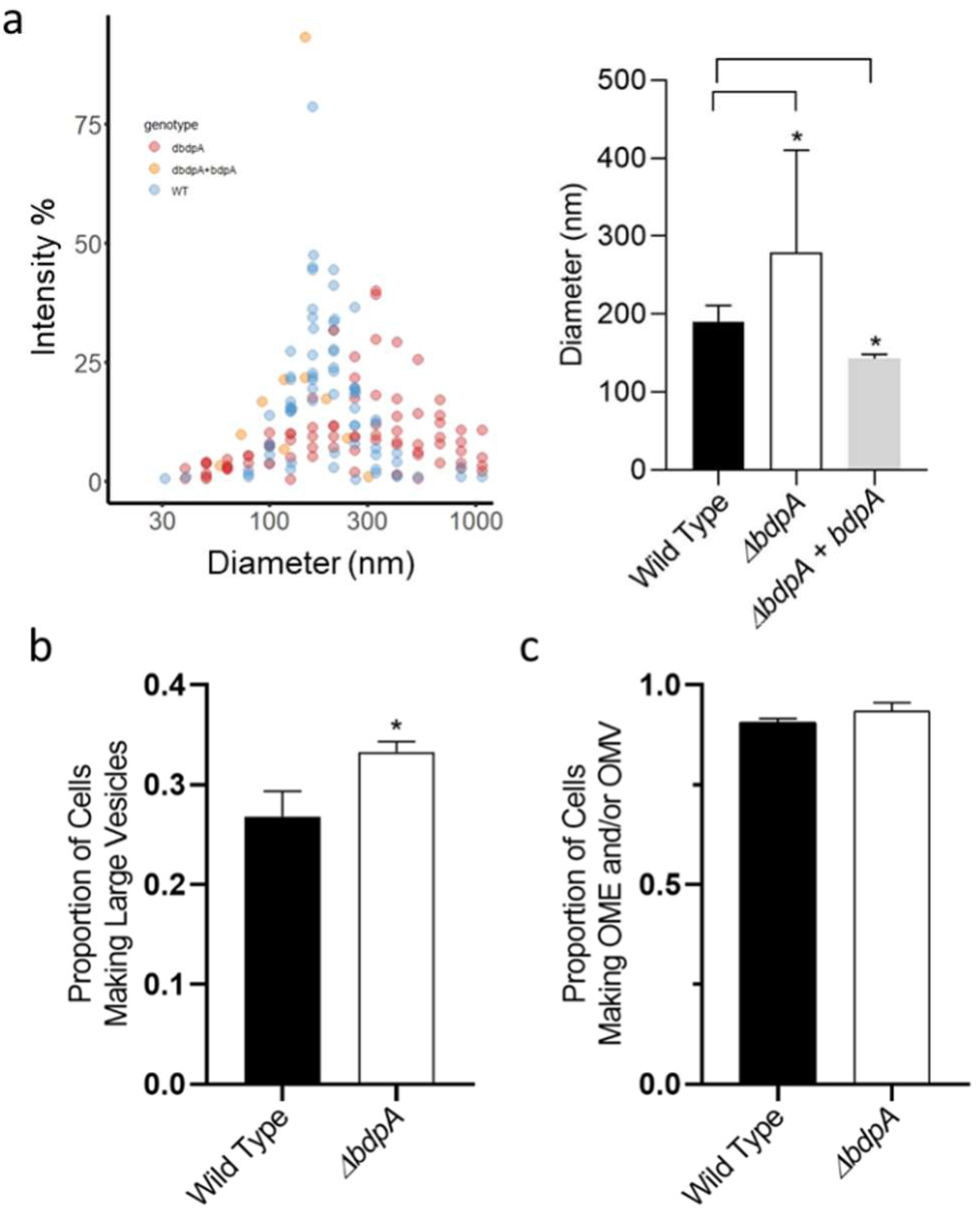
BdpA is responsible for maintaining vesicle size but does not alter the frequency of OMV or OME formation. **a**. Dynamic light scattering of OMV size distributions from the deletion strain (red, 9 biological replicates) compared to wild type (blue, 11 biological replicates) and Δ*bdpA* cells expressing *bdpA* from a plasmid (yellow, (left), with weighted averages of OMV diameters (right). Color opacity denotes overlapping data points (p < 0.05, *t*-test). **b**. Proportion of cells forming large vesicles (>300 nm diameter) during perfusion flow monitored by time lapse fluorescence imaging (p < 0.00459, χ^2^ test). **c**. Proportion of cells forming vesicles and extensions relative to the total number of cells observed by fluorescence microscopy during perfusion flow (p > 0.05, χ^2^ test). Time lapse images were recorded from 3 biological replicates per strain with 5-13 fields of view per replicate. Error bars represent standard deviation.

Cell-associated OMV frequency and size distribution was also measured in live cultures using a perfusion flow imaging platform and the membrane stain FM 4-64, as described previously (21). *S. oneidensis* strains were monitored for OME/V production and progression over the course of 5 hours (>5 fields of view per replicate, n=3) using time-lapse imaging. Spherical membrane stained extracellular structures were classified as OMVs, while larger aspect ratio (i.e. length greater than the width) structures were classified as OMEs. The proportion of cells producing ‘large’ vesicles, defined as having a membrane clearly delineated from the interior of the vesicles and typically >300 nm, was quantified, and Δ*bdpA* cells produced significantly more large vesicles compared to WT cells (Fig. 2b) even though both the overall frequency of vesiculation and extensions were the same (Fig. 2c). Previous studies showed that OMEs transition between large vesicles and OMEs over time (21). BdpA appears to be involved in this transition in *Shewanella* due to the increased frequency of large vesicles from Δ*bdpA* cells. The median diameter of the OMVs is also the apparent maximum diameter observed in outer membrane extensions (21) suggesting BdpA influences membrane morphologies of both structures.

### BdpA constrains membrane extension morphology

Cells were also visualized after deposition onto a glass coverslip instead of a perfusion flow chamber as previously reported (20). BdpA was expressed from a 2,4-diacetylphloroglucinol (DAPG)-inducible promoter (55) (P_PhlF_-BdpA) in the Δ*bdpA* strain containing the plasmid p452-*bdpA* with 12.5 μM DAPG. After 3 hours post deposition on cover glass, OMEs can be seen extending from WT, Δ*bdpA*, and Δ*bdpA* p452-*bdpA* cells (Fig. 3a, Supplemental Fig. 2, Supplemental videos 1-3, 5 fields of view, n=3). Similar to perfusion flow experiments, no statistically significant difference in the overall frequency of OME production was observed between the cells in static cultures. The resolution of fluorescence microscopy was insufficient to identify morphological differences between OMEs of wild type and mutant strains, therefore, cryo-TEM was used to assess OMEs in each of the strains at the ultrastructural level.

**Fig 3.**
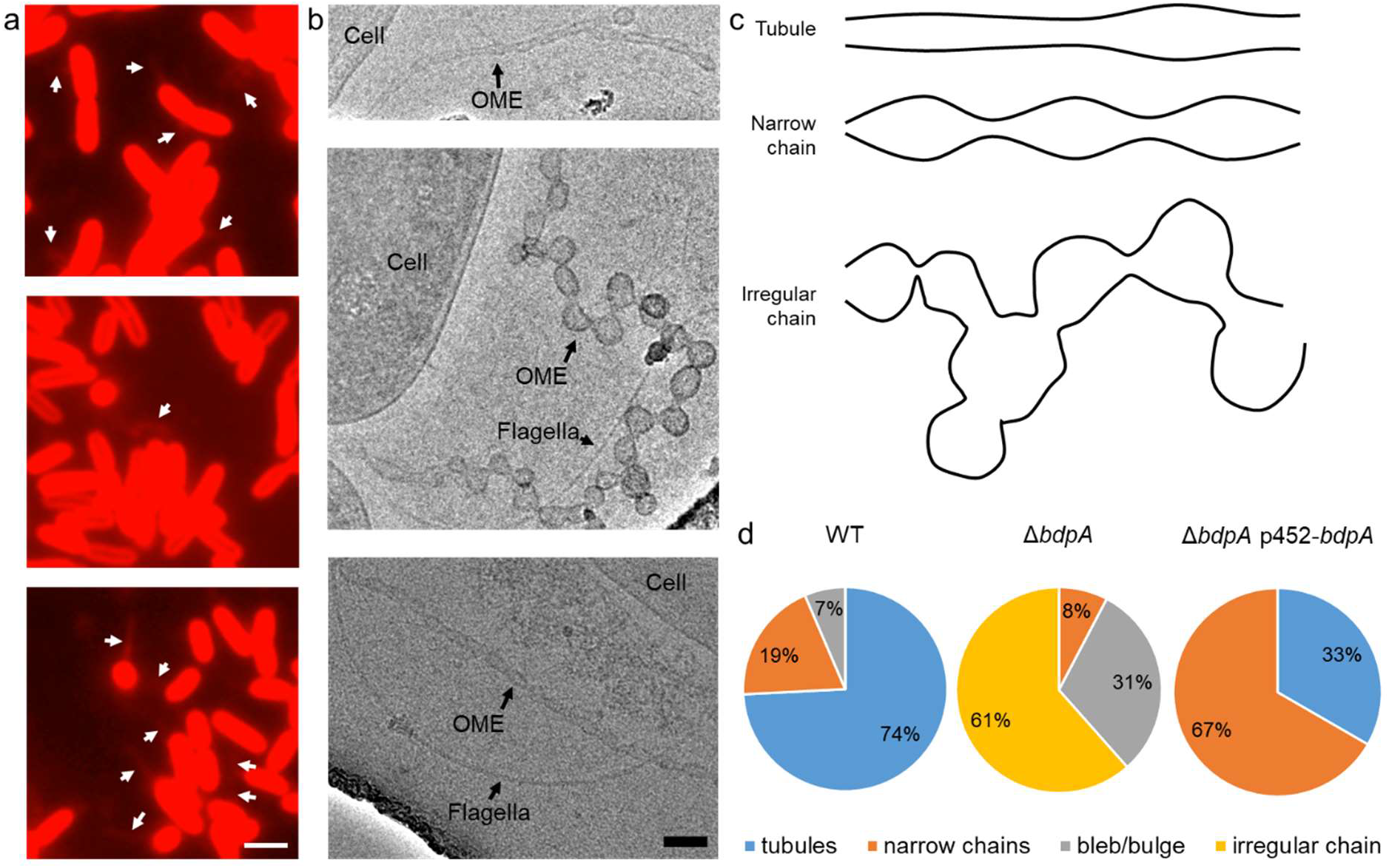
BdpA promotes OME maturation into ordered tubules. **a.** Fluorescence images of *S. oneidensis* WT (top), Δ*bdpA* (middle), and Δ*bdpA* p452-*bdpA* with 12.5 μM DAPG (bottom) OMEs. Scale = 2 μm. **b.** Cryo-TEM images of WT (top), Δ*bdpA* (middle), and Δ*bdpA* p452-*bdpA* with 12.5 μM DAPG (bottom) OMEs. Scale = 100 nm. **c.** Representative cartoon of OME phenotypes. **d**. Pie charts show relative frequency of OME phenotypes observed from each strain. Membrane blebs/bulges were defined as non-structured membrane protrusions that did not resemble either of the other OME categories.

*S. oneidensis* OMEs from unfixed WT, Δ*bdpA,* and Δ*bdpA* p452-*bdpA* strains were visualized at 90 minutes (Supplemental Fig. 3) and 3 hours (Fig. 3a) following cell deposition onto EM grids. By 3 hours post inoculation, images of WT cells consistently show narrow, tubule-like OMEs or vesicle chains of symmetric curvature (Fig. 3b,c n=31 OMEs observed). The Δ*bdpA* OMEs appear as lobed, disordered vesicle chains with irregular curvature compared to the WT (p < 0.001, Fisher’s exact test), and vesicles can be observed branching laterally from lobes on the extensions (Fig. 3b,c n=13 OMEs observed). WT OMEs also exhibited lateral branching of vesicles and lobes, but they exhibited uniform curvature and diameter between lobes, mirroring previous observations of nascent OMEs imaged immediately following OME formation (21). Tubules were not observed in any Δ*bdpA* OMEs at 3 hours (Fig. 3d). OMEs from Δ*bdpA* p452-*bdpA* cells appear as narrow tubules of a uniform curvature or as ordered vesicle chains (Fig. 3b,c n=3 OMEs observed), showing that expression of BdpA from a plasmid rescues the mutant phenotype by constricting and ordering OMEs into narrow tubules and chains (Fig. 3d, p < 0.01, Fisher’s exact test). Earlier in OME progression at 90 minutes, WT OME phenotypes appeared narrow, tubule-like, and seldom interspersed with lobed regions (Supplemental figure 3a). In Δ*bdpA* OMEs, lobed regions are prevalent with irregular curvature (Supplemental figure 3b). Several narrow Δ*bdpA* p452-*bdpA* OMEs evenly interspersed with slight constriction points or “junction densities” were observed extending from a single cell (Supplemental Fig. 3c).

### Expression of BdpA results in OMEs during planktonic growth

*S. oneidensis* OMEs are more commonly observed in surface attached cells than planktonic cells (20, 21). BAR domain proteins can directly promote tubule formation from liposomes *in vitro*(9), so inducing expression of an additional copy of the *bdpA* gene prior to attachment could result in OME formation even during planktonic growth. Growth curves were similar in cultures with the pBBR1-mcs2 empty vector in either of the WT or Δ*bdpA* background strains, but induction of *bdpA* in Δ*bdpA* p452-*bdpA* cells at higher concentrations of 1.25 and 12.5 μM 2,4-diacetylphloroglucinol affected the growth rate (Supplemental figure 4). Planktonic cultures inoculated from overnight cultures were induced with 12.5 μM DAPG for 1 hour, labeled with FM 4-64, and imaged by confocal microscopy. Neither WT (Fig. 4) nor WT with the empty plasmid exposed to 12.5 μM DAPG (not shown) displayed OMEs immediately following deposition onto cover glass. However, 12.5 μM DAPG-induced bdpA expression from p452-*bdpA* in the WT background strain (WT p452-*bdpA*) displayed OMEs immediately, ranging between 1-7 extensions per cell (Figure 4, Supplemental video 4). The abundance of OMEs suggest that increased *bdpA* expression in planktonic cultures can initiate membrane sculpting into OMEs at the expense of cell division.

**Fig 4.**
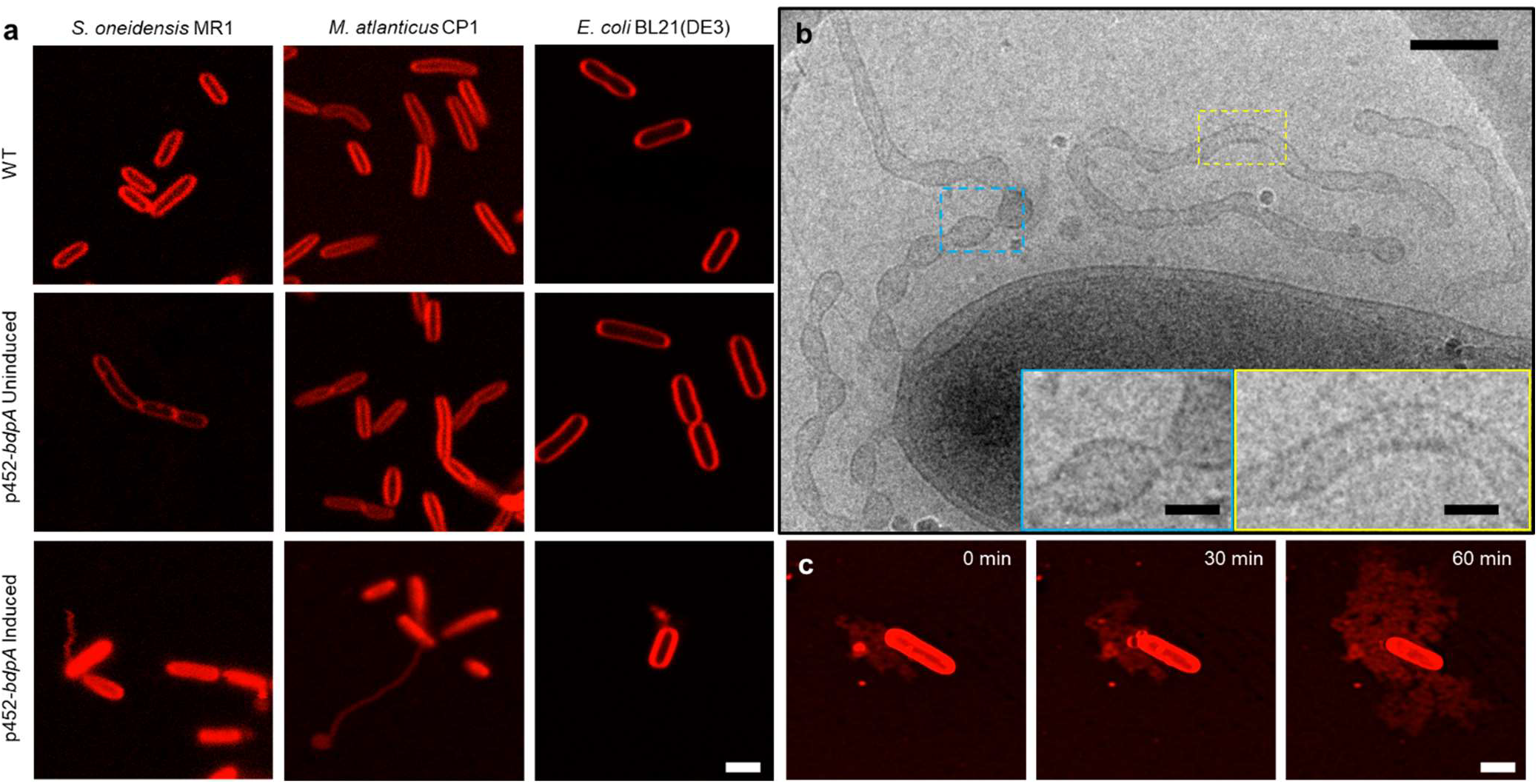
Heterologous expression of BdpA promotes OME formation. **a.**Induction of BdpA expression with 12.5 μM DAPG during planktonic, non-attached growth results in OME formation in *S. oneidensis* (left), *M. atlanticus* CP1 (middle), and *E. coli* BL21(DE3) (right). Scale = 2 μm. **b**. Cryo-TEM image of OMEs following planktonic induction of BdpA expression in *S. oneidensis* WT p452-*bdpA* cells. Scale = 200 nm. Insets enlarged to show detail of regularly ordered electron densities at the surface of OME junctions (blue) and tubule regions (yellow). Scale = 50 nm. **c.** OME growth over time at 30 minute intervals of *E. coli* BL21(DE3) expressing BdpA while attached to a glass surface. Scale = 2 μm.

The ultrastructure of OMEs resulting from expression of *bpdA* from WT p452-*bdpA* cells was examined by cryo-TEM, but in this case samples from planktonic cultures were vitrified on EM grids after induction rather than incubation during induction on the EM grids. By expressing extra copies of *bdpA* in the WT strain during planktonic induction, we predicted that OME morphology would be predominately reliant on curvature formation by BdpA rather than other unidentified structural proteins involved in intrinsic OME formation. OMEs appear as tubule-like segments interspersed with pearled regions proximal to the main cell body (Fig. 4b). OMEs from the MR-1 p452-*bdpA* strain are observed as thin, tubule-like outer membrane vesicle chains, suggesting BdpA involvement in the constriction of the larger outer membrane vesicle chains into longer, tubule-like extensions with more evenly interspersed junction densities. The BdpA OME phenotype more closely resembles membrane tubules formed by the F-BAR protein Pacsin1 from eukaryotic cells, showing a mixture of tubule regions interspersed with pearled segments (56, 57). Orientation and association of Pacsin1 dimers on the membrane surface with one another impacted tubule morphology, and BdpA membrane sculpting could be similar mechanistically.

### BdpA-mediated membrane extensions in *Marinobacter atlanticus* CP1 and *E. coli*

To test the effect of expressing BdpA in an organism with no predicted BAR domain-containing proteins and no apparent OME production, BdpA was expressed in *Marinobacter atlanticus* CP1 (58). *Marinobacter* and *Shewanella* are of the same phylogenetic order (*Alteromonadales*) and have been used for heterologous expression of other *S. oneidensis* proteins, such as MtrCAB (59, 60). Upon exposure to DAPG, *M. atlanticus* containing the p452-*bdpA* construct (CP1 p452-*bdpA*) forms membrane extensions (Figure 4). OMEs ranged from small membrane blebs to OME tubules extending up to greater than 10 μm in length from the surface of the cell (Supplemental Fig. 5). As noted previously, variation in the tubule phenotypes are commonly seen in tubules from eukaryotic F-BAR proteins (56, 57), showing possible mechanistic overlap of membrane curvature functionalities between these two separate BAR domain proteins.

**Fig 5.**
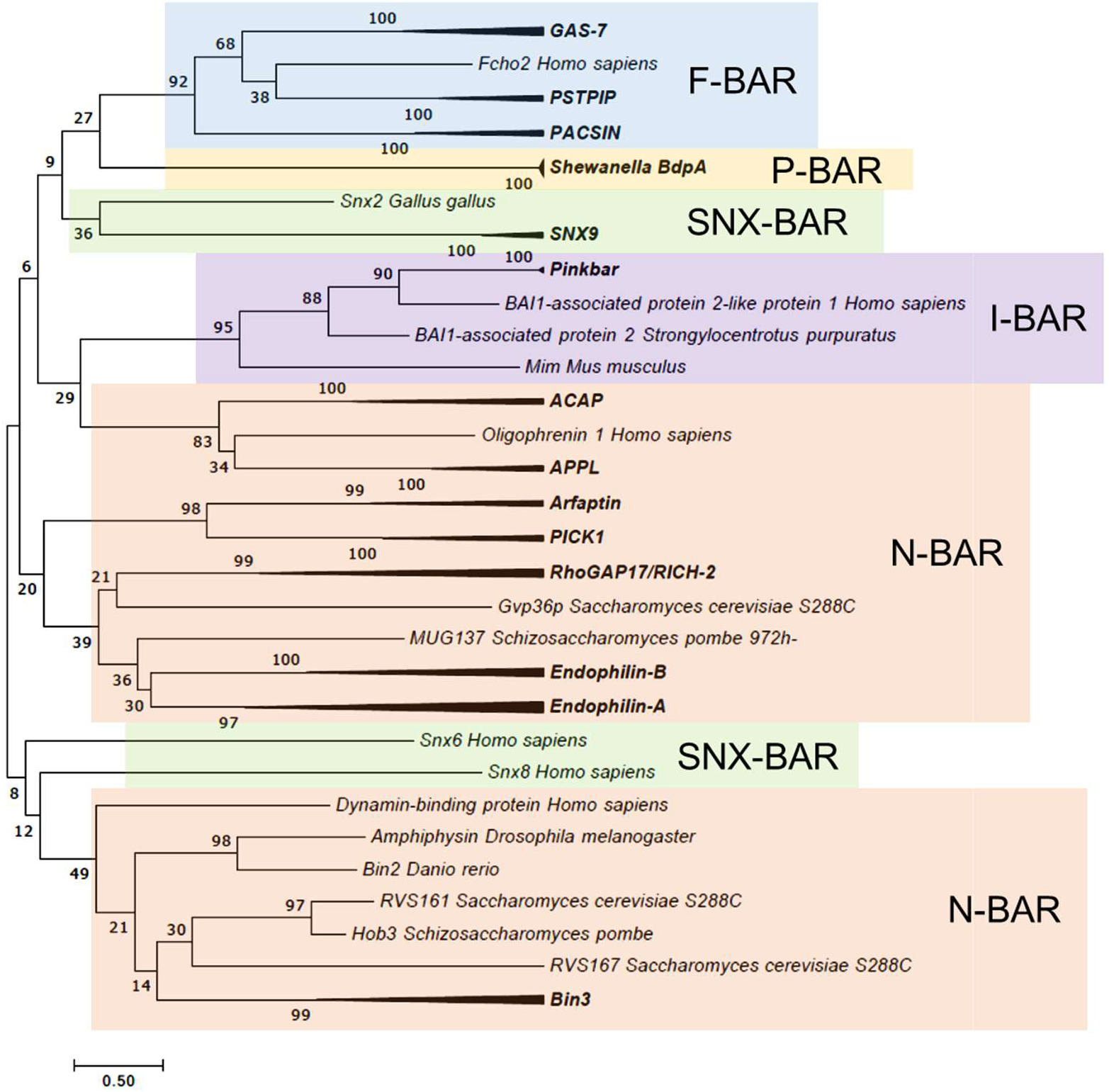
Comparative phylogenetic analysis of BdpA with prokaryotic homologs and eukaryotic BAR domains. Maximum Likelihood evolutionary histories were inferred from 1000 bootstrap replicates, and the percentage of trees in which the taxa clustered together is shown next to the branches. Arrows indicate multiple branches collapsed to a single node. *S. oneidensis* BdpA and 5 prokaryotic orthologs (WP_011623497 – unclassified *Shewanella*, ESE40074 – *S. decolorationis* S12, WP_039978560 - *S. decolorationis*, KEK29176 – *S. xiamenensis*, and WP_055648003 – *Shewanella sp.* Sh95) predicted by the current BAR domain Pfam HMM to contain a BAR domain aligned with representative BAR domains from various BAR domain subtypes (N-BAR, F-BAR, SNX-BAR, I-BAR) at a total of 196 positions. The Gamma distribution used to model evolutionary rate differences among sites was 12.9598.

In previous membrane curvature formation experiments with eukaryotic BAR domain proteins, localized BAR domain protein concentrations affected the resultant shape of the membranes, ranging from bulges to tubules and branched, reticular tubule networks at the highest protein densities (61–63). We predicted that expression of BdpA in cells optimized for protein overexpression, such *E. coli* BL21(DE3), would show OMEs resembling BAR protein concentration-dependent structures previously observed from eukaryotic BAR protein experiments *in vitro*. While the uninduced *E. coli* BL21(DE3) p452-*bdpA* cells had uniform, continuous cell membranes similar to those of plasmid-free BL21(DE3) cells under the conditions tested, *E. coli* BL21(DE3) cells containing the p452-*bdpA* vector induced with 12.5 μM DAPG had outer membrane extensions and vesicles (Figure 4). When visualized over time, OMEs progressed towards a network of reticular membrane structures extending from the cell (Fig. 4c). After 30 minutes, additional membrane blebs were observed that developed into elongated OMEs by 60 minutes. Growth of *E. coli* OMEs was coincident with shrinking of the cell body (from initial cell length = 4.5 μm to 3.5 μm at 60 minutes), supporting direct membrane sculpting activity of BdpA. *In vitro* tubule formation assays with purified proteins and liposomes are the canonical approach by which eukaryotic BAR domain proteins have been assessed for membrane sculpting activity. However, molecular crowding of purified proteins with no documented membrane curvature formation activity, such as GFP, can also lead to ordering of liposomes into tubules (64). Further, tubule formation from liposomes is not limited to BAR domain protein activity and requires non-physiologically high protein concentrations (65–67). The heterologous expression approach allows for more rapid screening of putative BAR domain proteins for membrane sculpting activity, avoiding the need for protein purification and *in vitro* systems.

### P-BAR: a new BAR domain subtype

The discovery of a novel, functional BAR domain protein in prokaryotes provokes questions into the evolutionary origin of BAR domains, such as whether the BdpA BAR domain in *S. oneidensis* arose as a result of convergent evolution, a horizontal gene transfer event, or has a last common ancestor across all domains of life. BdpA homologs were identified by PSI-BLAST in other *Gammaproteobacteria*, including most but not all species of *Shewanella*, as well as *Alishewanella*, *Rheinheimera*, and *Cellvibrio* (Supplemental Fig. 6). The current BAR domain Pfam Hidden Markov Model (HMM) prediction analysis identified BAR domain features in only 5 of the 52 prokaryotic homologs despite greater than 90% homology to *S. oneidensis* BdpA. An amino acid alignment of the 52 BdpA homologs was used to generate a maximum likelihood phylogenetic tree showing evolutionary relatedness of BdpA orthologs to the BAR domain prediction sequences (Supplemental Fig. 6). The 5 BdpA orthologs predicted to contain a BAR domain based on the current model were subsequently aligned with representative known BAR proteins from the various BAR domain subtypes (N-BAR, F-BAR, and I-BAR)(68). BdpA and its prokaryotic orthologs cluster separately from the eukaryotic BAR proteins in their own distinct clade (Fig. 5), suggesting that while BdpA contains a functional BAR domain, it represents its own class of BAR domain, hereafter named P-BAR (Prokaryotic BAR). The closest phylogenetic relative to P-BAR domains are the F-BAR domains, and BdpA membrane sculpting phenotypes are functionally similar to previous *in vitro* observations from Pacsin1 (56, 57). It is possible that the P-BAR domain arose as a result of horizontal gene transfer from a eukaryote due to the prevalence of eukaryotic coiled-coil proteins with predicted homology to BdpA after 2 iterations of PSI-BLAST. However, the branch lengths and low bootstrap values supporting the placement of P-BAR relative to other BAR domain subtypes make it challenging to directly infer the evolutionary history of P-BAR domains. Discovery of other putative P-BAR proteins would help to build this analysis, and if future comparative proteomics analysis of OME/Vs demonstrates overlapping activity of BdpA with preferential cargo loading into OME/Vs, it could hint at the evolutionary origins of vesicle-based protein trafficking. Conservation of BAR domain proteins supports the notion that three-dimensional organization of proteins in lipid structures is as important to prokaryotes as it is eukaryotes, and suggests additional novel P-BAR proteins are waiting to be discovered.

## Conclusion

*S. oneidensis* expresses a functional prokaryotic BAR domain protein, which is the first identified and characterized in bacteria. Enrichment of BdpA in the redox-active OMVs suggests overlapping mechanistic functionality with eukaryotic BAR proteins in the context of vesicle constriction (52). This finding was further demonstrated through fluorescence microscopy during perfusion flow, where large vesicles were more frequently observed from Δ*bdpA*. Membrane constriction activity of BdpA was confirmed through cryo-EM images that depicted an inability to transition into ordered tubules in the absence of *bdpA* expression. Variation in OMEs with BdpA ranged from ordered, narrow vesicle chains of a consistent diameter to stable tubules. The closest phylogenetic eukaryotic BAR domain subtype to BdpA, F-BAR domains, exhibit similar variation in tubule morphology, depending upon the orientation of the tip-to-tip oligomerization around the tubules (2, 3, 57). Subsequent studies will include vesicle constriction into tubules with purified protein to ascertain the extent of functional mechanistic similarity of BdpA to other F-BAR proteins. However, heterologous expression of BdpA and other potential P-BAR domain proteins enables rapid validation of membrane sculpting mechanistic activity. Ultimately, the discovery of BdpA and its homologs presents a critical first step in the new field of bacterial BAR domain protein research.

## Methods

### Bacterial strains, plasmids, and medium

The bacterial strains used in this study can be found in Supplemental Table 1. *S. oneidensis* strains were grown aerobically in Luria Bertani (LB) media at 30°C with 50 μg/mL kanamycin when maintaining the plasmid. To observe membrane extensions, cells were centrifuged and resuspended in a defined media comprised of 30 mM Pipes, 60 mM sodium DL-lactate as an electron donor, 28mM NH_4_Cl, 1.34 mM KCl, 4.35 mM NaH_2_PO_4_, 7.5 mM NaOH, 30 mM NaCl, 1mM MgCl_2_, 1mM CaCl_2_, and 0.05 mM ferric nitrilotriacetic acid(22). *Marinobacter atlanticus* CP1 strains were grown in BB media (50% LB media, 50% Marine broth) at 30°C with 100 μg/mL kanamycin to maintain the plasmids as described previously (58).

Inducible BdpA expression plasmids were constructed for use in *S. oneidensis* MR-1, *M. atlanticus* CP1, and *E. coli* BL21(DE3) using the pBBR1-mcs2 backbone described previously (58). The Marionette sensor components (*phlF* promoter, consitutively expressed PhlF repressor, and yellow fluorescence protein (YFP)) cassette from pAJM.452 (55) was cloned into the pBBR1-mcs2 backbone, and the YFP cassette was replaced with the gene encoding BdpA by Gibson assembly (primers in Supplemental Table 2). The resulting plasmid was given the name p452-*bdpA*. The Gibson assembly reactions were electroporated into *E. coli* Top10 DH5α cells (Invitrogen), and the sequences were confirmed through Sanger sequencing (Eurofins genomics). Plasmid constructs were chemically transformed into conjugation-competent *E. coli* WM3064 cells for conjugative transfer into the recipient bacterial strains of *S. oneidensis* MR-1 and *M. atlanticus* CP1. The same BdpA expression vector was transformed into *E. coli* BL21(DE3) cells (Invitrogen) by chemical transformation.

Generation of a scarless *ΔbdpA* knockout mutant of *S. oneidensis* was performed by combining 1 kilobase fragments flanking upstream and downstream from *bdpA* by Gibson assembly into the pSMV3 suicide vector. The resultant plasmid pSMV3_1507KO was transformed into *E. coli* DH5α λ*pir* strain UQ950 cells for propagation. Plasmid sequences were confirmed by Sanger sequencing before chemical transformation into *E. coli* WM3064 for conjugation into *S. oneidensis*. Conjugation of pSMV3_1507KO into *S. oneidensis* MR-1 was performed as described previously (23). Optical densities at 600nm were measured to determine growth curves for each strain in either LB or SDM with 50 μg/mL kanamycin and 2,4-diacetylphloroglucinol (DAPG) as indicated. Cultures of 500 μL samples diluted to an initial OD_600_ of 0.1 were grown in Costar® polystyrene 48 well plates (Corning Incorporated) within a Tecan Infinite M1000 Pro (Grödig, Austria) plate reader at 30°C with shaking agitation at 258 rpm. Optical densities were recorded every 15 minutes with i-Control software (2.7). All measurements were performed in triplicate.

### Purification of Outer Membrane Vesicles

*S. oneidensis* MR-1 cells were grown in LB in 1L non-baffled flasks at 30° C at 200 RPM. When an OD_600_ of 3.0 was reached, cells were pelleted by centrifugation at 5000 x g for 20 min at 4°C, resulting supernatant was filtered through a 0.45 μm filter to remove remaining bacterial cells. Vesicles were obtained by centrifugation at 38,400 x g for 1 h at 4°C in an Avanti J-20XP centrifuge (Beckman Coulter, Inc). Pelleted vesicles were resuspended in 20 ml of 50 mM HEPES (pH 6.8) and filtered through 0.22 μm pore size filters. Vesicles were again pelleted as described above and finally resuspended in 50 mM HEPES, pH 6.8, except for vesicle preparations used for electrochemistry which were suspended in 100 mM MES, 100 mM KCl, pH 6.8. Extracellular DNA, flagella, and pili can all be co-purified. Protocol was adapted from Perez-Cruz et al (69).

### Cryoelectron tomography

Vesicle samples were diluted to a protein concentration of 0.4 mg/mL and applied to glow-discharged, X-thick carbon-coated, R2/2, 200 mesh copper Quantifoil grid (Quantifoil Micro Tools) using a Vitrobot chamber (FEI). Grids were automatically plunge frozen and saved for subsequent imaging. No fixative was used. Images were collected on an FEI Krios transmission electron microscope equipped with a K2 Summit counting electron-detector camera (Gatan). Data were collected using customized scripts in SerialEM(70), with each tilt series ranging from −60° to 60° in 3° increments, an underfocus of ∼1-5 μm, and a cumulative electron dose of 121 e/A^2^ for each individual tilt series. Tomograms were reconstructed using a combination of ctffind4 (71) and the IMOD software package (72).

### Dynamic Light Scattering

Distributions of vesicle diameters were measured with Wyatt Technology’s Möbiuζ dynamic light scattering instrument with DYNAMICS software for data collection and analysis. Data was collected using a 0-50 mW laser at 830 nm. The scattered photons were detected at 90°. Measurements were recorded from 11 biological replicates for WT OMVs, and 9 replicates for Δ*bdpA* OMVs. Mobius software analyzed the population of particles to generate a table with binned diameters and the percentage of particles at each diameter. In order to compute the average vesicle size from each sample, a weighted average was computed so that diameter bins that had the most number of vesicles would be accurately represented in the final average weight of the population. The product of each diameter is multiplied by its percentage in the population, these products are added together for each sample, divided by the sum of weights. The weighted diameters per replicate were then averaged for each genotype. Statistical significance was determined by Student’s *t*-test, and error bars represent standard deviation.

### Electrochemistry

CHA Industries Mark 40 e-beam and thermal evaporator was used to deposit a 5 nm Ti adhesion layer and then a 100 nm Au layer onto cleaned glass coverslips (43mm x 50mm #1 Thermo Scientific Gold Seal Cover Glass, Portsmouth NH, USA). Self-assembled monolayers were formed by incubated the gold coverslip in a solution of 1mM 6-mercaptohexanoic acid in 200 proof ethanol for at least 2 hours. Electrode was then rinsed several time in ethanol followed by several rinses in milliQ water. The SAMs layer was then activated by incubation in 100 mM N-(3-Dimethylaminopropyl)-N′-ethylcarbodiimide hydrochloride and 25 mM N-hydroxysuccinimide, pH 4, for 30 minutes. A sample of outer membrane vesicles was deposited on the surface of the electrode and incubated at room temperature overnight in a humid environment. Cyclic voltammetry was performed in a 50 mL three-electrode half-cell completed with a platinum counter electrode, and a 1 M KCl Ag/AgCl reference electrode electrical controlled by a Gamry 600 potentiostat (Gamry, Warminster, PA). The whole experiment was completed in an anaerobic chamber with 95% nitrogen, 5% hydrogen atmosphere.

### Proteomics

Vesicle samples were prepared as described above. *S. oneidensis* outer membrane (OM) was purified via the Sarkosyl method described by Brown et al.(73). A 50 mL overnight culture of cells was harvested by centrifugation at 10,000 × g for 10 min. The cell pellet suspended in 20 mL of 20 mM ice-cold sodium phosphate (pH 7.5) and passed four times through a French Press (12000 lb/in^2^). The lysate was centrifuged at 5,000 × g for 30 min to remove unbroken cells. The remaining supernatant was centrifuged at 45,000 × g for 1 h to pellet membranes. Crude membranes were suspended in 20 mL 0.5% Sarkosyl in 20 mM sodium phosphate and shaken horizontally at 200 rpm for 30 min at room temperature. The crude membrane sample was centrifuged at 45,000 × g for 1 h to pellet the OM. The pellet of OM was washed in ice-cold sodium phosphate and recentrifuged.

To prepare for mass spectrometry samples were treated sequentially with urea, TCEP, iodoactinamide, lysl endopeptidase, trypsin, and formic acid. Peptides were then desalted by HPLC with a Microm Bioresources C8 peptide macrotrap (3×8mm). The digested samples were subjected to LC-MS/MS analysis on a nanoflow LC system, EASY-nLC 1200, (Thermo Fisher Scientific) coupled to a QExactive HF Orbitrap mass spectrometer (Thermo Fisher Scientific, Bremen, Germany) equipped with a Nanospray Flex ion source. Samples were directly loaded onto a PicoFrit column (New Objective, Woburn, MA) packed in house with ReproSil-Pur C18AQ 1.9 um resin (120A° pore size, Dr. Maisch, Ammerbuch, Germany). The 20 cm x 50 μm ID column was heated to 60° C. The peptides were separated with a 120 min gradient at a flow rate of 220 nL/min. The gradient was as follows: 2–6% Solvent B (7.5 min), 6-25% B (82.5 min), and 25-40% B (30 min) and to 100% B (9min). Solvent A consisted of 97.8% H2O, 2% acetonitrile, and 0.2% formic acid and solvent B consisted of 19.8% H2O, 80% ACN, and 0.2% formic acid. The QExactive HF Orbitrap was operated in data dependent mode with the Tune (version 2.7 SP1build 2659) instrument control software. Spray voltage was set to 2.5 kV, S-lens RF level at 50, and heated capillary at 275 °C. Full scan resolution was set to 60,000 at m/z 200. Full scan target was 3 × 106 with a maximum injection time of 15 ms. Mass range was set to 300−1650 m/z. For data dependent MS2 scans the loop count was 12, target value was set at 1 × 105, and intensity threshold was kept at 1 × 105. Isolation width was set at 1.2 m/z and a fixed first mass of 100 was used. Normalized collision energy was set at 28. Peptide match was set to off, and isotope exclusion was on. Data acquisition was controlled by Xcalibur (4.0.27.13) and all data was acquired in profile mode.

### Bioinformatics

Putative BAR domain SO_1507 (BdpA) was identified in search of annotation terms of *S. oneidensis* MR-1. The conserved domain database (CDD-search)(NCBI) was accessed to identify the position-specific scoring matrix (PSSM) of the specific region of SO_1507 that represented the BAR domain (amino acid residues at positions 276-421). The domain prediction matched to BAR superfamily cl12013 and specifically to the family member BAR cd07307. LOGICOIL multi-state coiled-coil oligomeric state prediction was used to predict the presence of coiled-coils within BdpA (43). SignalP 6.1 was used to detect the presence of the signal peptide and cellular localization of BdpA (74). I-TASSER protein structure prediction (47) was used to generate a predicted model of the BAR domain region of BdpA. Alignment of the predicted BdpA BAR domain structure model to the structure of Hof1p was performed in the PyMOL Molecular Graphics System, Version 2.0 (Schrödinger, LLC).

A PSI-BLAST (75) search against the NCBI nr database was performed using the BdpA BAR sequence as the initial search seed to determine how prevalent the BdpA BAR domain is in related species. Conserved BdpA orthologs were annotated as hypothetical proteins in all of the species identified. In the initial round, 24 proteins were found from other organisms identified as *Shewanella* with a high conservation among the proteins and another 28 proteins were found in more distant bacteria species that had similarity of 65% to 44%. A second iteration identified a few proteins much more distantly related from bacterial species and then proteins from eukaryote phylum *Arthropoda* that were annotated as being centrosomal proteins. All of the found proteins from bacterial species were hypothetical proteins with no known function. Only five of the proteins from the search returned hits to the PSSM of the BAR cd07307. The identity among the proteins was very high and examination of the proteins suggests that a functional form similar to the BAR domain would result for all the found proteins. Overall, this places BdpA as a protein that just barely meets criteria via PSSM models to be assigned as matching the BAR domain while the rest of the proteins found have enough differences to fail to match the BAR model while still being very similar to BdpA. An attempt was made to build up a HMM (Hidden Markov Model) using hmmer (76) to use for searching for other proteins that might match, but as with the PSI-BLAST search, only the proteins that formed the model returned as good matches. There appears to be a tight clade of very similar proteins with very little differentiation in the sequence. This indicates that while sequence homology between BdpA and the existing BAR domain consensus sequence predicted the BAR domain region in BdpA using hmmer or NCBI tools, the sequence conservation is at the cusp of a positive hit by the HMM since other closely related (>90% homology) BdpA orthologs were not predicted to contain a BAR domain by this method. The most homologous eukaryotic protein to BdpA (27%) is a putative centrosomal protein in *Vollenhovia emeryi* (accession #: XP_011868153) that is predicted to contain an amino terminal C2 membrane binding domain and a carboxy-terminal SMC domain within a coiled-coil region. Despite CDD search failing to predict the presence of a BAR domain in this protein, it does not preclude the presence of one, pending an updated BAR Pfam HMM. Alignments and phylogenies were constructed in MEGA 7. MUSCLE was used to align protein sequences, and Maximum Likelihood phylogenies were inferred using the Le-Gascuel (LG+G) substitution matrix (77). Initial trees for the heuristic search were obtained automatically by applying Neighbor-Join and BioNJ algorithms to a matrix of pairwise distances estimated using a JTT model, and then selecting the topology with superior log likelihood value. A discrete Gamma distribution was used to model evolutionary rate differences among sites as indicated in figure legends. All positions with less than 85% site coverage were eliminated.

### Confocal microscopy

For *in vivo* imaging of intrinsic outer membrane extension production, *S. oneidensis* MR-1 strains were grown in LB media overnight, washed twice with SDM, and diluted to an OD_600_ of 0.05 in 1 mL of SDM with appropriate antibiotics. Prior to pipetting, ~1cm of the pipette tip was trimmed to minimize shear forces during transfer. 100 μL of each culture was labeled with 1 μL 1M FM 4-64 to visualize the cell membranes. After staining, 10 μL of the labeled cell suspension was gently pipetted onto 22 x 22 mm No.1 cover glass (VWR) and sealed onto glass slides with clear acrylic nail polish for confocal imaging or onto Lab-Tek chambered #1 cover glass (Thermo Fischer Scientific)(for widefield fluorescence). On average, intrinsic membrane extension formation could be observed starting after 45 minutes sealed onto cover glass. For planktonic OME production from BdpA induction in trans, diluted cells (in either SDM for *S. oneidensis*, BB for *M. atlanticus*, or LB for *E. coli*) were induced with 12.5 μM DAPG for 1 hour at 30°C with 200 RPM shaking agitation. Cells were labeled with FM 4-64 and sealed onto glass slides as before. Induced OMEs were imaged immediately after mounting onto slides.

Confocal images were taken by a Zeiss LSM 800 confocal microscope with a Plan-Apochromat 63x/1.4 numerical aperture oil immersion M27 objective. FM 4-64 fluorescence was excited at 506 nm: 0.20% laser power. Emission spectra was detected from 592-700 nm using the LSM 800 GaAsP-Pmt2 detector. To capture the dynamics of the OMEs, images were collected over the designated length of time between 0.27 – 0.63 seconds per frame. Single frame time series images were collected of either a 50.71 μm by 50.71 μm (2x zoom) or a 20.28 μm by 20.28 μm (5x zoom) field of view. Widefield fluorescence images were taken using a LED-Module 511 nm light source at 74.2% intensity with 583-600 nm filters and a 91 He CFP/YFP/mCherry reflector. Excitation and emission spectra were 506 nm and 751 nm, respectively. Images were collected using a Hamamatsu camera with a 250 ms exposure time. Images were recorded using the Zeiss Zen software (Carl Zeiss Microscopy, LLC, Thornwood, NY, USA). Frequency of OMEs was calculated from 3 biological replicates of each strain from at least 5 fields of view selected at random (2444 cells from WT, 4378 from Δ*bdpA*, and 3354 from Δ*bdpA* p452-*bdpA*). OMEs were counted per cell per field of view from a 20 second video if a membrane extension was observed extending from the cell over that timeframe. Statistical significance was determined by two-tailed Student’s *t*-test.

### Perfusion flow microscopy

For OME statistics comparing *S. oneidensis* strains MR-1 and Δ*bdpA,* cells were pre-grown aerobically from frozen (−80°C) stock in 10 mL of Luria-Bertani (LB) broth (supplemented with 50 μg/mL Kanamycin for strains with plasmid) in a 125-mL flask overnight at 30°C and 225 rpm. The next day, the stationary phase (OD_600_ 3.0 – 3.3) preculture was used to inoculate 1:100 into 10 mL of fresh LB medium in a 125-mL flask. After ~6 hours at 30°C and 225 rpm, when the OD_600_ was 2.4 (late log phase), 5 mL of cells were collected by centrifugation at 4226 x *g* for 5 min and washed twice in defined medium. The perfusion chamber, microscope, and flow medium described previously (20–22) were used for all perfusion flow OME statistics experiments. During each 5 hour imaging experiment, the perfusion chamber was first filled with this flow medium, then <1 mL of washed cells were slowly injected for a surface density of ~100-300 cells per 112 x 112 μm field of view on a Nikon Eclipse Ti-E inverted microscope with the NIS-Elements AR software. Cells were allowed to attach for 5-15 minutes on the coverslip before perfusion flow was resumed at a volumetric flow rate of 6.25 ± 0.1 μL/s. Cells and OMEs were visualized with the red membrane stain FM 4-64FX in the flow medium (0.25 μg/mL of flow medium). A total of 1,831 wild type and 2,265 Δ*bdpA* cells were used for extension and vesicle quantification. Experiments were performed from three individual biological replicates for each strain, and statistical significance was determined by two-tailed χ^2^ test.

### Cryo transmission electron microscopy

*Shewanella* strains were streaked onto LB plates with or without kanamycin and allowed to incubate 3 days on a benchtop. The night before freezing, individual colonies were inoculated into 3 ml LB +/− kanamycin and incubated at 30 °C overnight with 200 rpm shaking. The following morning optical densities of the cultures were measured at 600nm and adjusted to a final OD_600_ of 1. Cells were pelleted at 8,000 rpm for three minutes for buffer exchange/washes. For the Δ*bdpA* p452-*bdpA* transformed cells, 12.5 μM DAPG was added. A freshly glow discharged 200 mesh copper grid with R2/1 Quantifoil carbon film was placed into a concavity slide. Approximately 150 μl of a 1:10 dilution of the cell suspensions, with or without the inducer, was added to cover the grid. A glass coverslip was then lowered onto the concavity to exclude air bubbles. The edges of the coverslip were then sealed with nail polish to prevent media evaporation. The slide assembly was then incubated in a 30 °C incubator for 1.5 to 3 hours. Immediately prior to plunge freezing, the top coverslip was removed by scoring the nail polish with a razor blade. TEM grids with cells were gently retrieved with forceps and loaded into a Leica grid plunge for automated blotting and plunging into LN2-cooled liquid ethane. Vitrified grids were transferred to a LN2 storage dewar. Imaging of frozen samples was performed on either a Titan (ThermoFisher Scientific) microscope equipped with a Gatan Ultrascan camera and operating at 300 kV or a Talos (ThermoFisher Scientific) equipped with a Ceta camera and operating at 200 kV. Images were acquired at 10,000x to 20,000x magnification and were adjusted by bandpass filtering. Unfixed OMEs were sorted based on appearance into 4 categories. Tubules were narrow OMEs with relatively uniform or slight symmetric curvature.

Narrow chains were recorded as OMEs with a narrow, consistent diameter and symmetric curvature at constriction points. Irregular chains were classified as OMEs without a consistent diameter throughout the length of the OME and asymmetric curvature on either side of the extension. Blebs \ bulges were outer membrane structures that did not resemble OMEs but still extended from the cell membrane surface. Phenotypes were documented from observations of 31 WT, 13 Δ*bdpA*, and 3 Δ*bdpA* p452-*bdpA* OMEs over three separate biological replicates, with two technical replicates of each strain per biological replicate. Two-tailed statistical significance between strains was calculated by Fisher’s exact test with the Freeman-Halton extension for a 2×4 matrix (78) using the Real Statistics Resource Pack software (Release 6.8).

## Acknowledgements

We thank Dr. Jeffery Gralnick for helpful discussions and advice; Dr. Adam Meyer and Dr. Chris Voigt for the DAPG-inducible Marionette promoter; Dr. Annie Moradian and Dr. Mike Sweredoski and the California Institute of Technology Proteome Exploration Lab for useful discussions on the preparation and analysis of proteomics data. Some of the cryo-TEM work was done in the Beckman Institute Resource Center for Transmission Electron Microscopy at Caltech. This work was supported by the United States Department of Defense Synthetic Biology for Military Environments (SBME) Applied Research for the Advancement of Science and Technology Priorities (ARAP) program. Work in ME-N’s lab was supported by the U.S. Office of Naval Research Multidisciplinary University Research Initiative Grant No. N00014-18-1-2632. LAZ was partially supported by the National Science Foundation grant DEB-1542527. SX was supported by the Division of Chemical Sciences, Geosciences, and Biosciences, Office of Basic Energy Sciences of the U.S. Department of Energy through grant DE-FG02-13ER16415. Work in GJJ’s lab was supported by the National Institute of Health (GM122588 to GJJ).

## Author Contributions

DP and LZ conceived the study independently then combined projects when complementary data on BdpA was discovered. LZ purified OMVs, prepared samples for LC MS-MS, and performed DLS measurements. LZ and SX made electrochemical measurements and analysis. DP conducted BdpA domain prediction and validation analysis, generated the p452-*bdpA* plasmid, Δ*bdpA* and p452-*bdpA* strains. DP and GC conducted fluorescence imaging experiments, and DP, LZ, and GC analyzed the data. LB adapted the Marionette sensor (PphlF-YFP) into pBBR1-mcs2. LZ and LAM performed cryo-TEM of OMVs. GC and LZ analyzed perfusion flow system data. CH, DP, and LD performed cryo-TEM experiments of OMEs and image processing / analysis. DP and AM generated phylogenetic data, and DP, AM, and BE analyzed the data. DP, LZ, CM, GC, AM, LAM, BE, GJJ, LD, MEN, and SG provided data interpretation. DP, LZ, MEN, and SG wrote the manuscript, with input from all coauthors.

---

**Supplemental Table 2.**
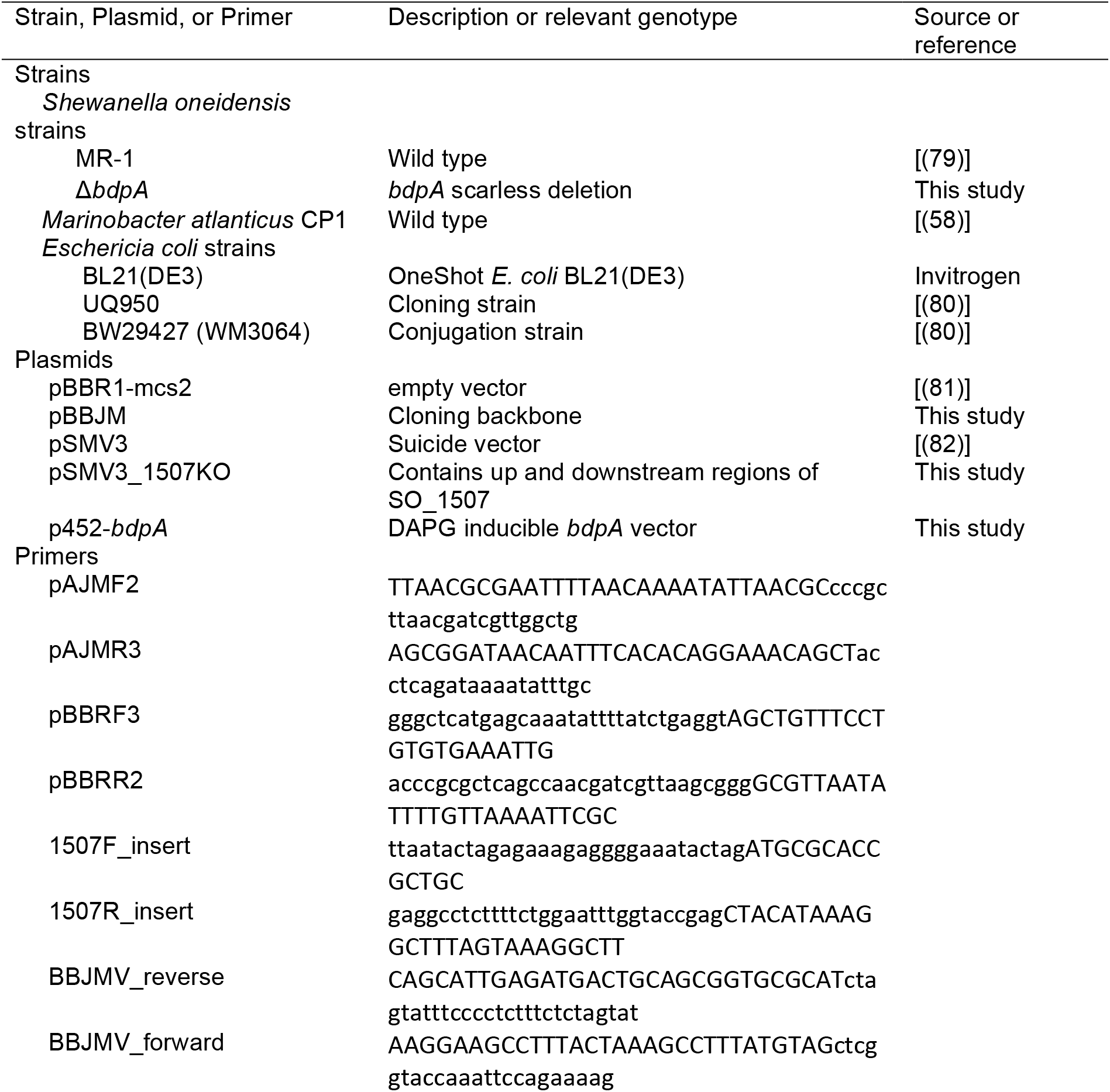

**Supplementary Figure 1.**
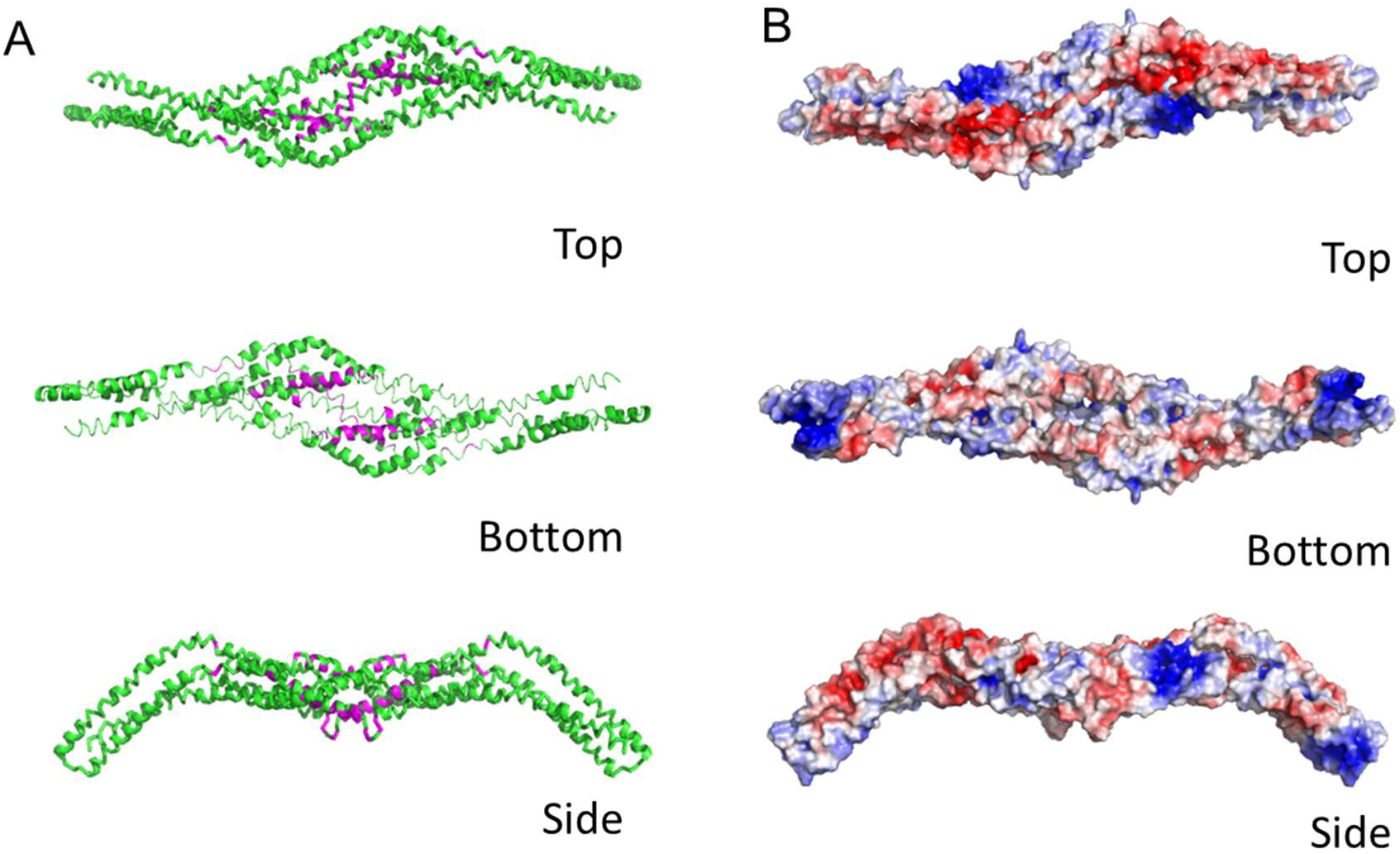
Predicted structure of the **BAR** domain region of **BdpA**. (A) Ribbon diagram of i-TASSER prediction of BdpA structure as a dimeric molecule, residues 175-502. Dimer was modeled from an alignment of BdpA monomers to homodimeric Hof1p (Protein Data Bank [PDB] ID code 4WPE) structures that resulted in closest proximity of putative dimer interface (purple) residues from the initial BAR domain prediction. (B) Surface representation of predicted BdpA homodimer colored according to electrostatic potential. The bottom concave face has an accumulation of distributed positively charged residues (Lys, Arg, His - blue), while the top face has clusters of positive and negatively charged (Glu, Asp - red) residues, within a range of −3.0 to 3.0 V.

**Supplementary Figure 2.**
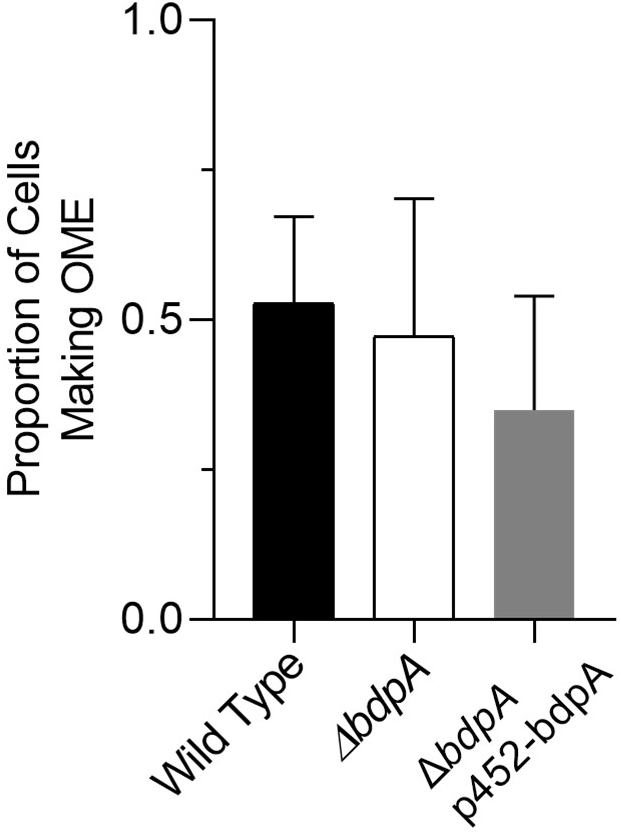
Proportion of cells making OMEs without perfusion flow at three hours post deposition onto chambered cover glass, measured from five random fields of view from each of three independent cultures per strain.

**Supplementary Figure 3.**
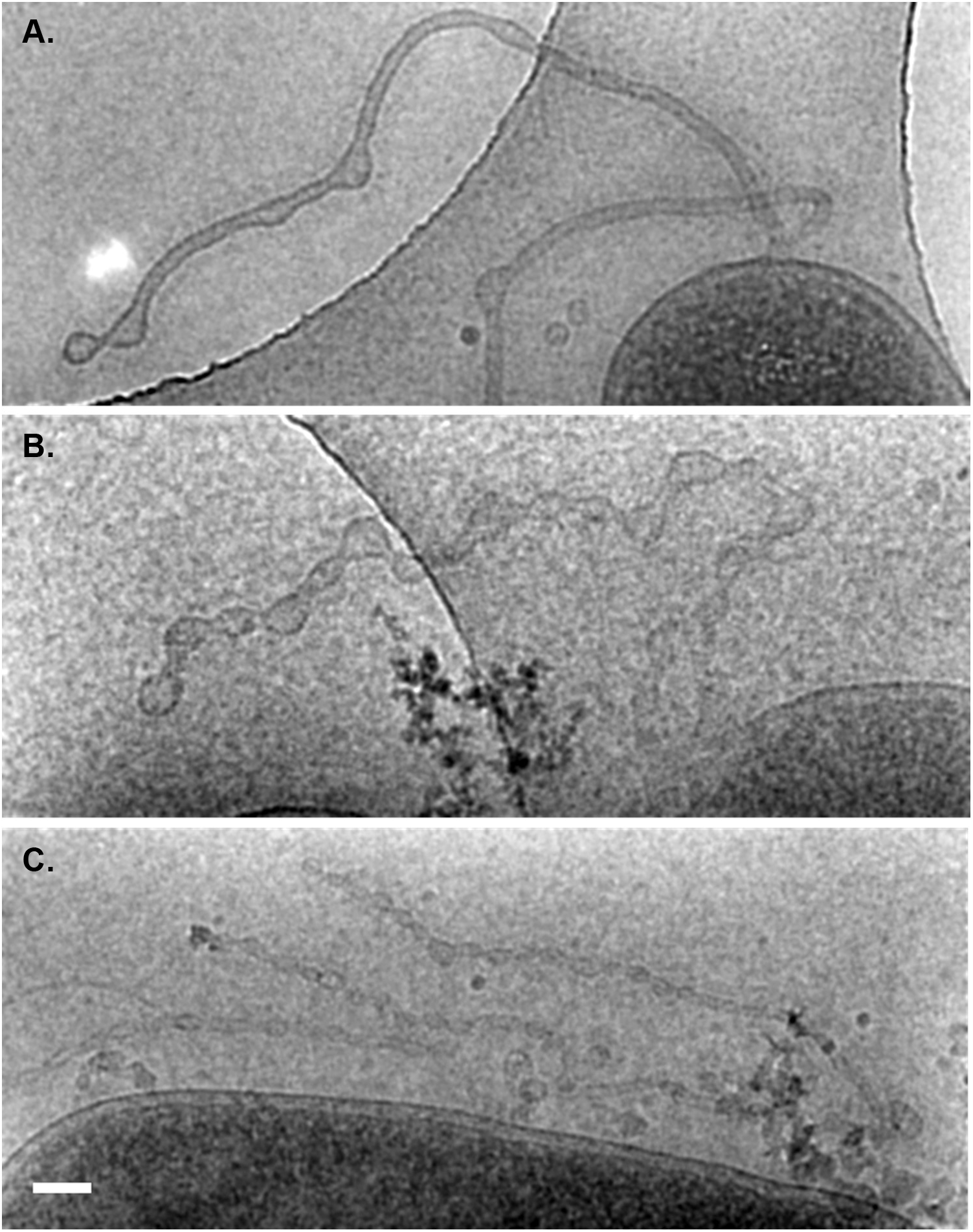
Cryo-TEM of *S. oneidensis* WT (A), Δ*bdpA* (B), and Δ*bdpA* p452-*bdpA* (C) OMEs at 90 minutes post-surface attachment. Scale = 100 nm.

**Supplementary Figure 4.**
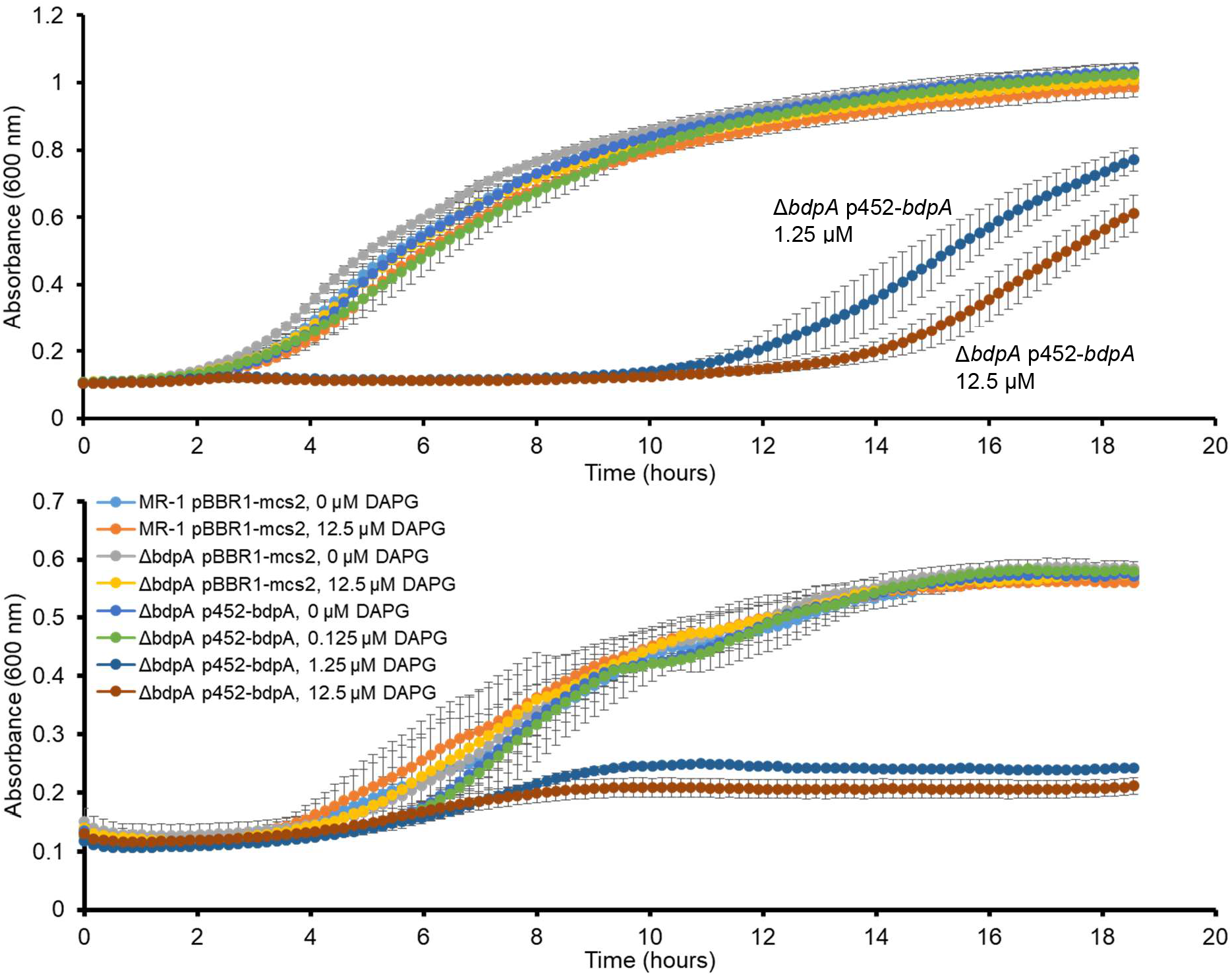
Growth of *S. oneidensis* strains in LB (top) or *Shewanella* Defined Medium (SDM) (bottom) in response to DAPG exposure and BdpA induction. Error bars are standard deviation of three biological replicates.

**Supplementary Figure 5.**
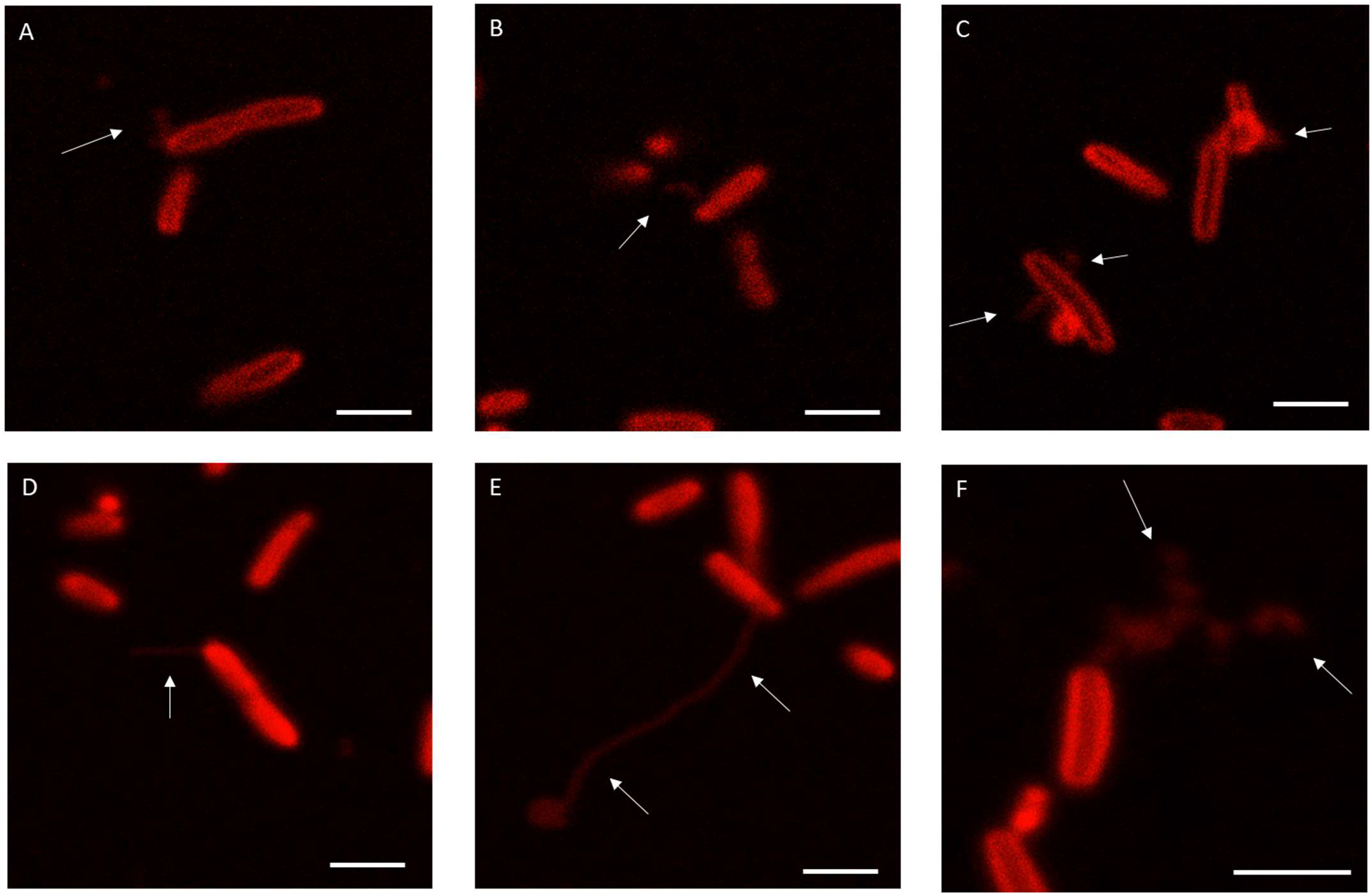
Variability in OME phenotypes following BdpA induction in *M. atlanticus* CP1 p452-*bdpA* cells. Cells displayed an array of membrane curvature phenotypes, ranging from short OMEs (<2μm, A-C), long OMEs (2-10+ μm, D-F), membrane blebbing (C,F), and branched OME/OMV chains (F). Scale = 2 μm.

**Supplementary Figure 6.**
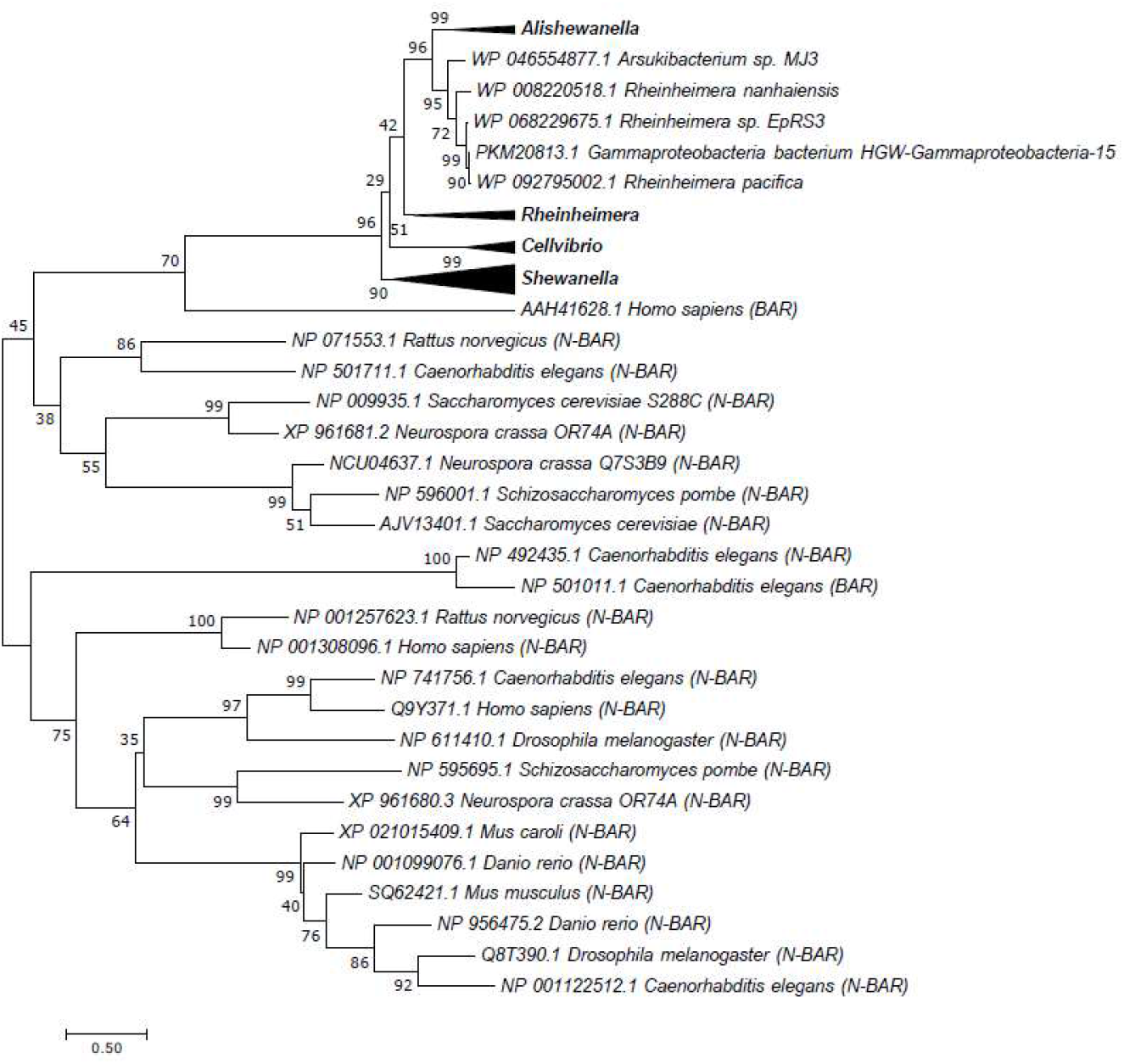
BdpA has homologs in other bacterial species. A phylogenetic tree of the 23 BAR domain sequences that seed the BAR domain HMM predictions, *S. oneidensis* BdpA, and conserved BdpA orthologs in other prokaryotes. The protein regions corresponding to the BdpA BAR domain sequence from the 52 prokaryotic BdpA orthologs were aligned with the 23 representative eukaryotic BAR domain-containing proteins used to generate the BAR domain consensus sequence (cd07307) at a total of 222 positions. Maximum Likelihood evolutionary histories were inferred from 1000 bootstrap replicates, and the percentage of trees in which the taxa clustered together is shown next to the branches. The Gamma distribution used to model evolutionary rate differences among sites was 11.9548.

**Supplementary Video 1** Widefield imaging of *S. oneidensis* WT cells 3h post-deposition onto the surface of a chambered cover glass. Scale = 5 μm.

**Supplementary Video 2** Widefield imaging of *S. oneidensis* Δ*bdpA* cells 3h post-deposition onto the surface of a chambered cover glass. Scale = 5 μm.

**Supplementary Video 3** Widefield imaging of *S. oneidensis* Δ*bdpA* p452-*bdpA* cells 3h post-deposition onto the surface of a chambered cover glass. Scale = 5 μm.

**Supplementary Video 4** Confocal imaging of *S. oneidensis* MR-1 p452-*bdpA* cells after 1h planktonic induction of BdpA with 12.5 μM DAPG. Scale = 5 μm.

